# Know your RNA-Seq data in depth: a case study using data from early life stress in mouse

**DOI:** 10.1101/2025.06.30.654452

**Authors:** Angelica Lindlöf

**Affiliations:** School of Bioscience, Systems Biology Research Centre, University of Skövde, Sweden

## Abstract

Next-generation sequencing (NGS) is a technology that enables rapid and high-throughput sequencing of entire genomes, transcriptomes or specific DNA/RNA populations. RNA-Seq is an NGS-based method that specifically targets the transcriptome and can be applied to bulk tissue or single cells. NGS produces large volumes of partial sequences (reads), which must be aligned, assembled and analyzed to extract meaningful biological information such as gene expression and genetic variants. However, NGS data often contain noise and errors due to technical factors like PCR bias, contamination or alignment inaccuracies. Understanding and managing this noise is important for ensuring the reliability of results, especially in clinical or diagnostic contexts. Quality control is a critical step in the data analysis process to ensure the accuracy, reliability and reproducibility of sequencing outcomes. In this study, a detailed quality assessment of RNA-Seq data is presented using a publicly available dataset from Usui et al. (2021). Read alignment was performed with the BWA-MEM2 tool. Quality control included analysis of reports generated by the FastQC and MultiQC tools, followed by in-depth examination of information contained in the resulting SAM/BAM files. Specifically, read alignments were evaluated for the FLAG status of paired reads, variant information extracted from CIGAR and MD strings, Mapped and Matched Identity metrics, chromosomal distribution of mapped reads and nucleotide-level mapping. This comprehensive analysis highlights the importance of variant profiling and alignment quality metrics in ensuring the reliability of RNA-Seq data.

## Introduction

Next-generation sequencing (NGS) is an experimental technology that enables relatively rapid, high-throughput sequencing of entire genomes, transcriptomes or specific DNA/RNA populations (Akintunde et al., 2025; McCombie et al., 2019). The key steps in NGS include fragmentation of DNA/RNA molecules into smaller pieces, attachment of adapters to the ends of these fragments that allows them to be captured and amplified, and sequencing of the amplified fragments using a selected NGS platform. These amplified fragments are only partially sequenced at one end (single-end reads) or both ends (paired-end reads), with the length of the partial sequences (read length) determined by the experimental protocol. Commonly chosen read length is 50-150 base pairs (bp) for short-read sequencing, whereas for long-read sequencing it is typically 10-30 kb bp. RNA-Seq is an NGS-based method that specifically targets the transcriptome, i.e., the complete set of RNA transcripts that have been extracted from a sample or tissue (bulk RNA-Seq) or a single cell (single-cell RNA-Seq).

NGS produces large volumes of raw data in the form of reads, which must undergo extensive processing and analysis (McCombie et al., 2019; Reuter et al., 2015). This generally involves aligning reads to a reference genome (if available), assembling them into longer sequences and subsequently identifying genetic variants, quantifying gene expression, or uncovering other biological features of interest. Moreover, NGS data inherently contains a level of noise, which is defined as inaccuracies or unwanted variations in the reads that arise during the sequencing process and are not due to biological factors such as natural selection. These errors can affect data interpretation and may lead to incorrect conclusions. Common sources of error and noise include incorrect base calling, PCR-induced base misincorporations, amplification and fragmentation bias, adapter or sample contamination, instrumental variability, sample quality issues, read alignment errors, variant calling errors, and biases in the reference genome.

Having an in-depth knowledge of the level of variants in your data are important for implementing strategies to handle their impact and ensure reliable sequencing results. High-quality data is essential for drawing accurate biological conclusions and forms the foundation for informed decision-making, particularly in clinical and diagnostic settings. Quality control of NGS data is one analysis step to ensure the accuracy, reliability and reproducibility of sequencing results.

In this study, we explore various metrics to thoroughly assess RNA-Seq data using a publicly available dataset from Usui et al. (2021); this research investigated the effects of early life stress (ELS) on the prefrontal cortex (PFC), a brain region critical for social behavior and emotional regulation. The researchers aimed to understand how ELS influences gene expression and the structural organization of the PFC, and how these changes correlate with social behavior and anxiety levels. Mice were used as the animal model to simulate ELS by subjecting weaned pre-adolescent mice to social isolation until adolescence. RNA-Seq was performed on three control mice and three ELS-exposed mice at 9 weeks of age, using RNA extracted from the prefrontal cortex. Sequencing was conducted with 151 bp paired-end reads on the Illumina NextSeq 500 platform (Usui et al., 2021).

The quality analysis includes examination of reports generated by the FastQC tool and summarized using the MultiQC tool, which effectively consolidates the reports from the two FastQ files generated during paired-end sequencing (Andrews, 2010; Cock et al., 2010; Ewels et al., 2016). Read alignment was performed using the BWA-MEM2 tool and the resulting SAM/BAM files were subsequently analyzed with respect to (Li & Durbin, 2010; Li et al., 2009):

1. The FLAG status of properly paired reads as reported in SAM/BAM files,
2. Variant information extracted from the CIGAR and MD strings reported in SAM/BAM files,
3. Mapped Identity and Matched Identity metrics and their role in filtering low-quality alignments,
4. The chromosomal distribution of mapped reads and
5. Nucleotide-level mapping results.

### FastQC/MultiQC analysis

The primary output of NGS is the raw sequencing reads, which are most commonly stored in FASTQ files. These are text-based files that contain both the nucleotide sequences and their corresponding sequencing quality scores in a standardized format called FASTQ (Cock et al., 2010). In the file, each read is represented by four lines referring to 1) a sequence identifier, 2) the actual sequence, 3) a separator line and 4) quality scores corresponding to each base in the sequence. The quality scores are typically in the form Phred scores, which are numerical values that represent the accuracy of each nucleotide base call in a read; a higher Phred score indicates greater confidence that the correct base has been called. General guidelines suggest that a score of ≥30 for at least 80-90% of the bases is indicative of high-quality sequencing data.

FastQC is a widely used bioinformatics tool for quality control of high-throughput sequencing data (Andrews, 2010). The tool provides a variety of analyses to help researchers assess the quality of sequencing reads generated in NGS experiments. In paired-end sequencing, two FASTQ files are generated per sample, one for each read direction. These paired reads are typically referred to as the forward (read 1) and reverse (read 2) reads. When using FastQC with paired-end data, a separate report is generated for each FASTQ file. MultiQC is a complementary tool designed to aggregate and summarize outputs from multiple bioinformatics analyses, and is commonly used to combine and present the two FastQC reports for paired-end data in a single, comprehensive summary (Ewels et al., 2016). In this study, we will examine three of the analyses reported by FastQC and summarized by MultiQC. These analyses will be illustrated using results from two representative samples, ELS3 and CTL3, from the mouse study by Usui et al. (2021). While these examples are highlighted, it is worth noting that the quality metrics are similar across all samples in the dataset.

The *Per Base Sequence Quality* plot displays the mean Phred quality score at each base position across all reads (Figure 1). In high-quality data, most scores should exceed Q30, indicating 99.9% base call accuracy, and fall within the green zone of the plot. For both samples, all base positions remain within this green area, with the lowest mean value still above 30, suggesting overall high-quality data. As commonly observed in NGS datasets, the quality scores are slightly lower at the first ∼10 bases, gradually decline toward the end of the reads, and tend to be somewhat lower for reverse reads.

**Figure 1.**
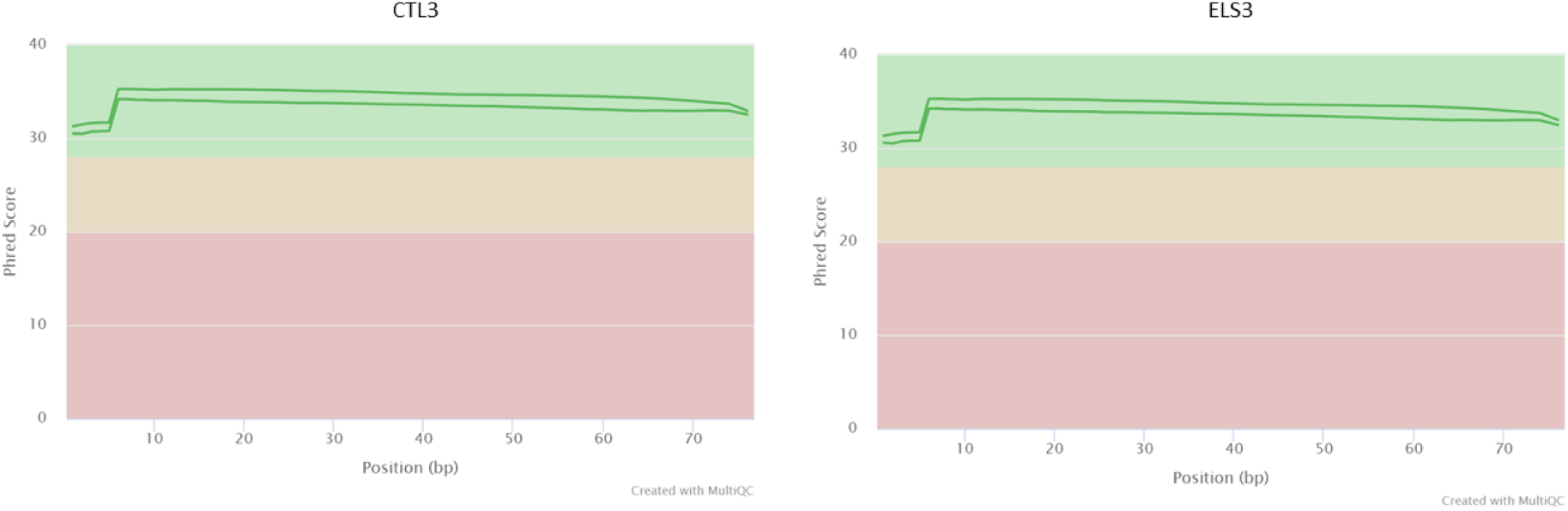
Per Base Sequence Quality plots generated with FastQC and summarized with MultiQC for samples CTL3 (left) and ELS3 (right) in the data from Usui et al. (2021). The x-axis represents base position in the read and y-axis the mean Phread quality score across the base position.

The *Per Sequence Quality Scores* plot shows the distribution of average quality scores across all reads (Figure 2). In high-quality datasets, most reads should have average scores of ≥30. For both samples, the majority of reads fall within the green area of the plot, indicating that their average quality scores meet this high-quality threshold. As is typical, reverse reads generally show slightly lower scores. A small proportion of reads in both samples exhibit low quality, falling within the yellow and pink areas. Additionally, the ELS3 sample shows slightly lower overall scores compared to CTL3, but the difference is not alarmingly lower.

**Figure 2.**
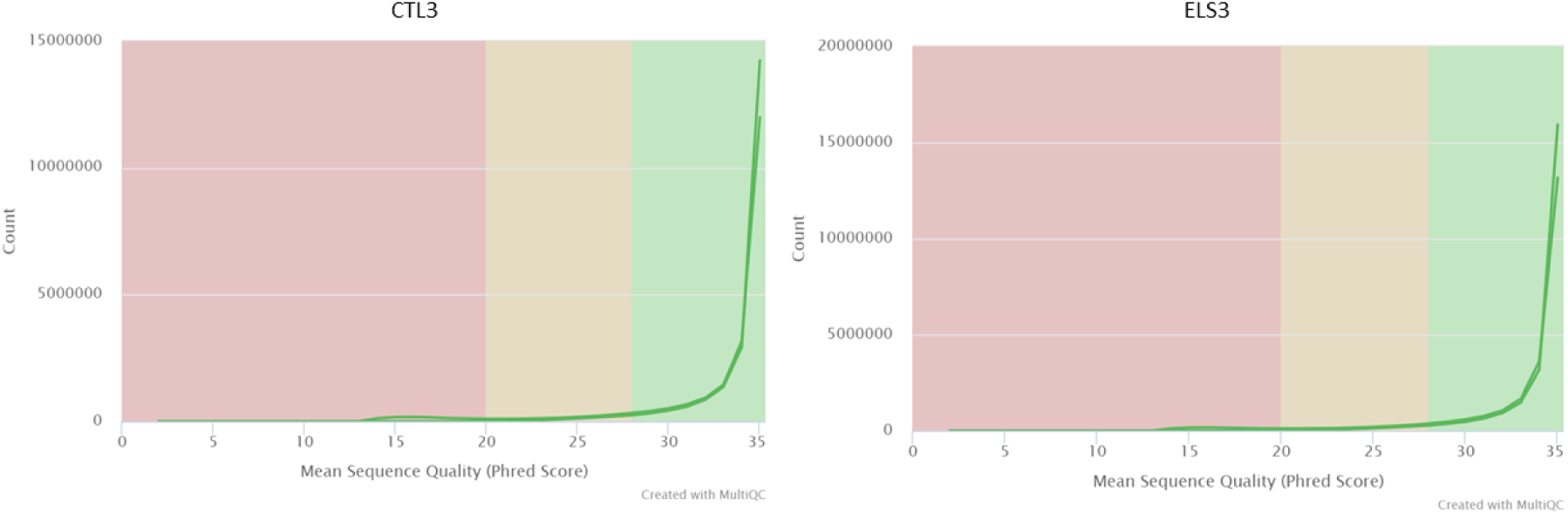
Per Sequence Quality Scores plots generated with FastQC and summarized with MultiQC for samples CTL3 (left) and ELS3 (right) in the data from Usui et al. (2021). The x-axis represents the average Phred quality score for a read and y-axis the number of reads having that score.

The *Sequence Length Distribution* plot shows the distribution of read lengths, with consistent read lengths expected in high-quality data (Figure 3). For both samples, the read lengths are uniform, without irregular bumps or peaks, and the majority of reads exceed 70 base pairs. Notably, the read lengths in the ELS3 sample are slightly longer, which likely corresponds to the slightly lower Per Sequence Quality Scores observed in this sample (see Figure 2 for comparison), as sequence quality typically decreases with increasing read length.

**Figure 3.**
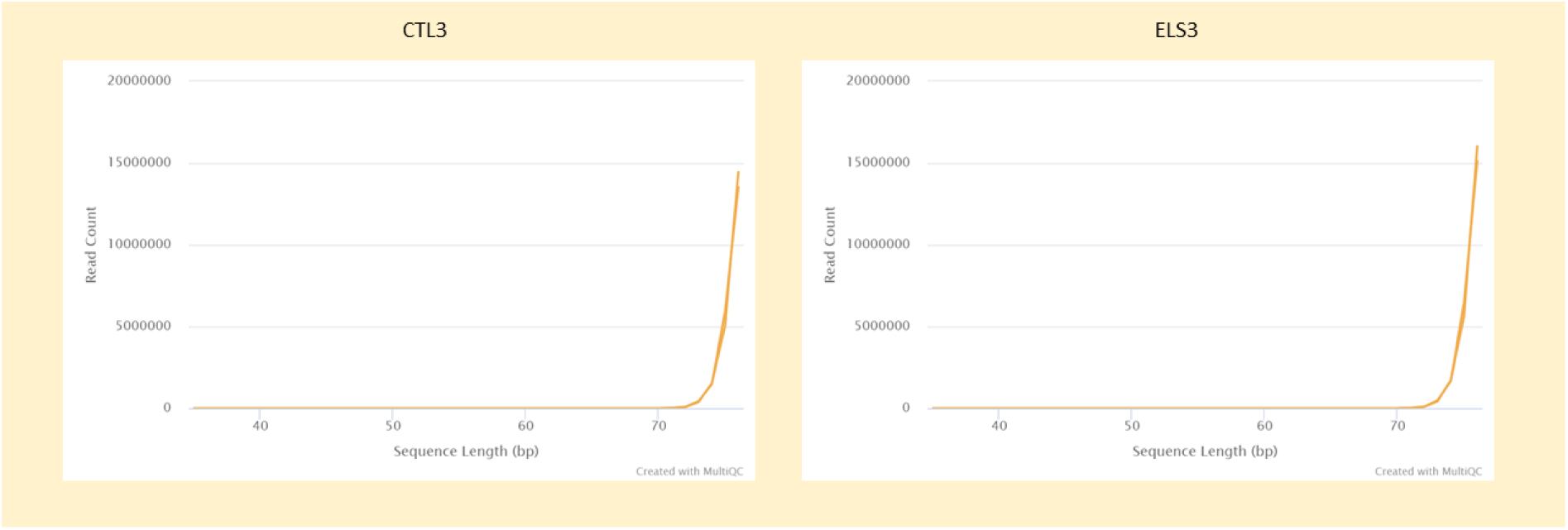
Sequence Length Distribution plots generated with FastQC and summarized with MultiQC for samples CTL3 (left) and ELS3 (right) in the data from Usui et al. (2021). The x-axis represents sequence length in bp and y-axis the number of reads having that length.

Overall, this dataset is considered high-quality, as evidenced by consistently high Phred quality scores and uniform read lengths. The majority of reads not only meet the quality threshold but also maintain satisfactory lengths, supporting reliable downstream analyses and accurate biological interpretations.

### Read alignment

Read alignment is an important step in the analysis of NGS data and comprises the mapping of sequencing reads back to a reference genome or transcriptome to identify their genomic origin. This step is foundational for many downstream analyses, including variant calling, gene expression quantification and detection of structural variants. Numerous alignment tools are available today, each differing in key aspects such as algorithmic approach, computational performance and suitability for specific types of DNA or RNA populations. Selecting the appropriate aligner depends on the characteristics of the dataset and the goals of the study.

In this study, BWA-MEM2 was chosen for read alignment (Li & Durbin, 2010). This tool supports alignment of both long and short reads, as well as single-end and paired-end reads, to a reference genome. The algorithm implemented in the tool is based on backward search using the Burrows-Wheeler Transform (BWT), allowing for mismatches and gaps in alignments. BWA-MEM2 is optimized for speed and memory efficiency, making it well-suited for large-scale genomic datasets and produces alignments with high accuracy. The tool was run with default settings, except that the output alignments were sorted by read names rather than by chromosomal position.

BWA-MEM2 also performs soft clipping during the alignment process, which helps manage reads that partially match the reference genome. When a segment of a read does not align well, BWA-MEM2 “soft clips” these bases, meaning that the clipped bases are not included in the alignment but are still retained in the read data. This approach helps remove sequencing errors commonly found at read ends without discarding entire reads. Additionally, soft clipping can remove adapter sequences or other contaminants present at the read ends. If reads are of high quality and have minimal adapter contamination or low-quality ends, trimming prior to alignment is often unnecessary, as soft clipping effectively handles these issues. In this study, since the dataset was deemed high-quality and only a small fraction (∼2%) of reads showed adapter contamination in the FastQC/MultiQC reports (data not shown), trimming was considered unnecessary before running BWA-MEM2.

The input for BWA-MEM2 is the FASTQ file generated from sequencing and the result is one or several alignments for each read. The number of alignments depends on whether a read is uniquely mapped (aligned to a single chromosomal location) or multi-mapped (aligned to multiple locations in the reference genome). These alignments are stored in Sequence Alignment/Map (SAM) files, with one SAM file generated per sample (Li et al., 2009). For paired-end data, alignments for both reads are contained within the same SAM file. Because SAM files can be very large, they are commonly converted to Binary Alignment Map (BAM) files, which is a compressed binary format of the SAM file but stores the information more efficiently.

The BAM files generated were subsequently analyzed using custom Python scripts. The parsing of BAM file lines was performed with the `bamnostic` package and utilizing the implementation developed by Chon and Huang (2021) for multi-processing BAM files (Chon & Huang, 2021; Sherman & Mills, 2018).

### SAM FLAGs

The SAM is a text-based file format developed by Li et al. (2009) to store information about sequences aligned to a reference genome and contains the aligned sequences along with various mapping details as well as alignment quality metrics. The file begins with a header section that provides metadata about the reference sequences, read and sample information, optional processing steps and comments. Following the header, each alignment is represented by a row consisting of 11 mandatory fields and a variable number of optional fields.

One of the mandatory fields in a SAM file is the FLAG, which is an integer value that encodes various attributes of a read’s mapping. These attributes include whether the read was successfully mapped to the reference, whether it is part of a paired-end read, or whether the alignment is primary or secondary. In a perfect paired-end sequencing (Figure 4), the two reads from a pair should align to the reference genome such that one read maps to the forward strand and the other to the reverse strand. Additionally, the reads should face each other, meaning the first read aligns to the forward strand and the second read to the reverse strand. The distance between the paired reads, known as the insert size, should fall within the expected range based on the library preparation. Finally, both reads should align to the same reference sequence, such as the same chromosome or contig.

**Figure 4.**
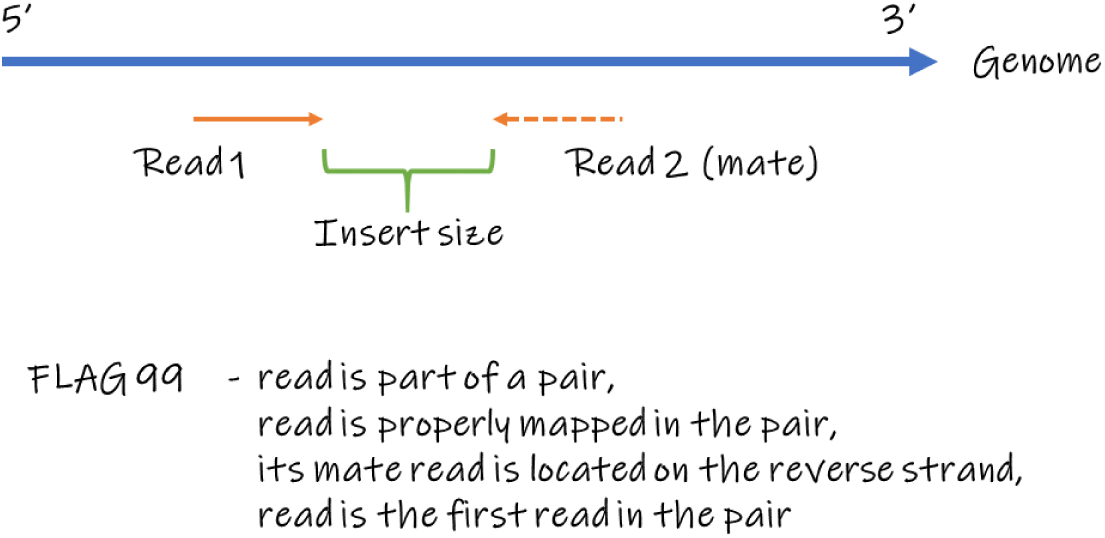
This schematic illustrates paired-end read alignment to a reference genome, showing Read 1 and Read 2 (aka mate). Read 1 aligns to the forward strand (5’ to 3’ direction), while Read 2 aligns to the reverse strand (3’ to 5’ direction), facing each other. The distance between the two reads on the genome is referred to as the insert size. The FLAG value 99 indicates that the read is part of a properly paired alignment where Read 1 is mapped, Read 2 is located on the reverse strand and Read 1 is the first in the pair.

The following FLAG values represent that a read in a pair could be properly aligned to the reference in a way that is consistent with the expected insert size, orientation and reference location, and means:

- 99 - the read is part of a pair, is properly mapped in the pair, its mate read (i.e., the second read sequenced from the fragment) is located on the reverse strand, and is the first read in the pair,
- 83 - the read is part of a pair, is properly mapped in the pair, is mapped to the reverse strand and is the first read in the pair,
- 163 - the read is part of a pair, is properly mapped in the pair, its mate read is located on the reverse strand, and is the second read in the pair,
- 147 - the read is part of a pair, is properly mapped in the pair, is mapped to the reverse strand and is the second read in the pair.

Reads assigned with any of these flags are referred to as properly paired.

### FLAG analysis

For the case study data, the alignment results show that a similar number of reads were sequenced across all samples, with sample ELS2 having the highest read count and CTL2 the lowest (Table 1). The proportion of properly paired reads was consistent across samples, at approximately 79%.

**Table 1.** Statistics regarding total number of reads (Total; counting reads sequenced from both ends) sequenced in each sample and the proportion of reads having the flag 99, 83, 163 or 147 (Properly paired) respectively reads having any other flag value (Other).

|  | CTL1 | CTL2 | CTL3 | ELS1 | ELS2 | ELS3 |
| --- | --- | --- | --- | --- | --- | --- |
| Total | 46,894,541 | 43,859,473 | 44,733,495 | 50,107,279 | 54,711,209 | 49,670,950 |
| Properly paired | 78.5% | 78.9% | 78.9% | 78.6% | 79.0% | 79.0% |
| Other | 21.5% | 21.1% | 21.1% | 21.4% | 21.0% | 21.0% |

Regarding reads that were assigned with any of the other FLAG values, the most common set were 81, 161, 97 and 145, which refers to reads that could not be properly mapped in the pair, i.e., either one of the reads is mapped in the wrong strand orientation, the distance between the two reads does not fall within the expected range or the reads do not align to the same reference sequence (Figure 5).

**Figure 5.**
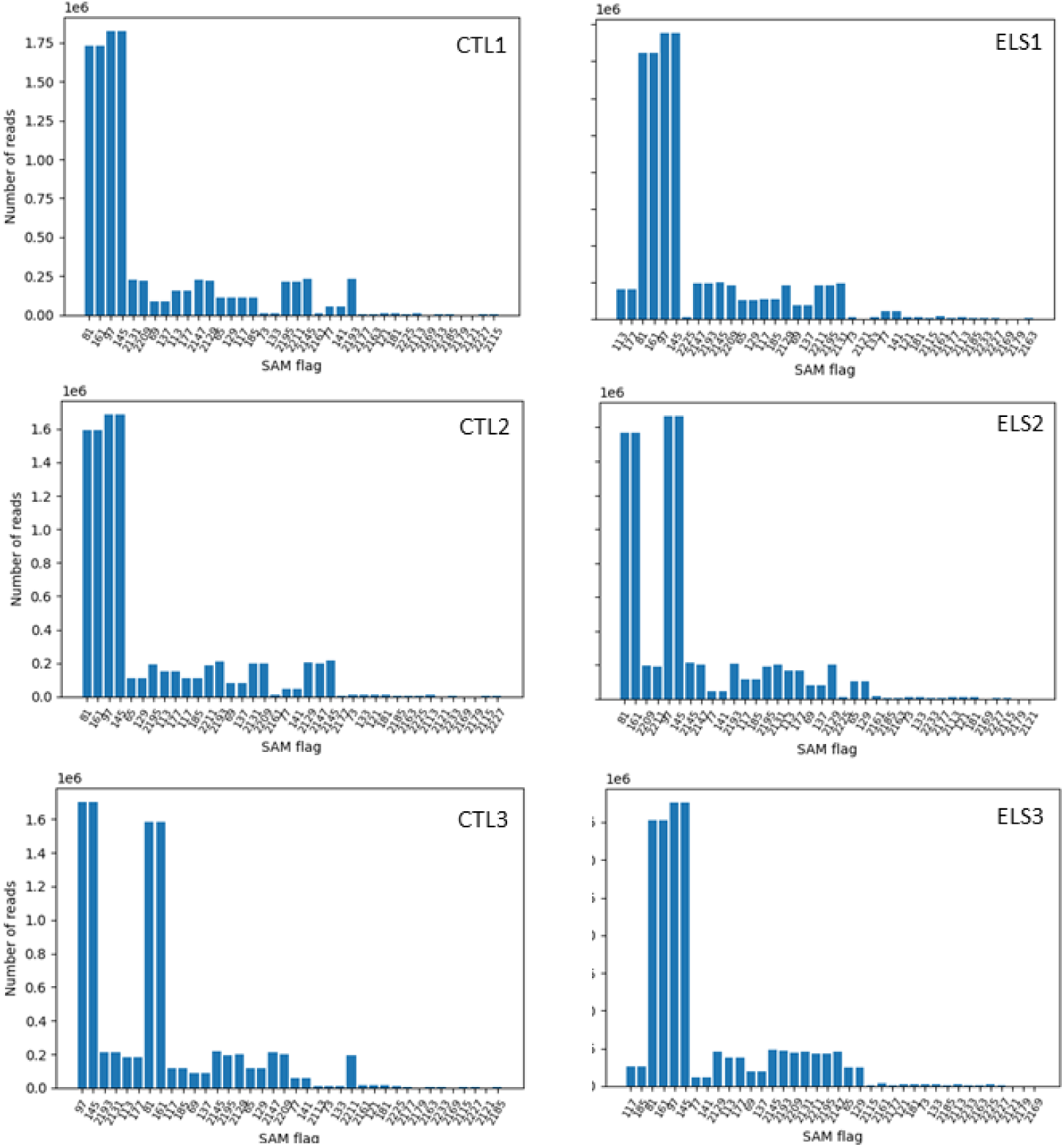
Histogram plots showing the number of reads per SAM FLAG value in each sample. The histogram does not include the FLAG values for properly paired paired-end reads. The x-axis refers to different SAM FLAG values and the y-axis number of reads assigned with one of the FLAG values.

Properly paired reads are critical for accurate interpretation of paired-end sequencing data and provide confidence that the reads are correctly aligned and thereby the data is reliable. Improper pairing could be due to errors during sequencing, chimeric reads, structural variations or mapping to low complexity regions. Such reads should be handled separately and could be analyzed for specific purposes, e.g., identification of structural variants, fusion genes or chimeric transcripts. About 21% of the reads in the samples could not be properly paired and *these reads were discarded from subsequent analyses*.

### BWA-MEM2 CIGAR and MD

The mandatory SAM file field Compact Idiosyncratic Gapped Alignment Report (CIGAR) provides detailed information on how each base in a read aligns to the reference genome. It describes various alignment operations such as matches, insertions, deletions and soft-clipped bases. Each operation is represented by a specific letter; for example, ‘M’ for alignment match, ‘S’ for soft clipping, ‘D’ for deletion and ‘I’ for insertion. These letters are preceded by an integer indicating the number of consecutive bases involved in the operation. For instance, ‘3M’ means three consecutive bases are matched to the reference, while ‘2M5D2M1I’ indicates 2 matched bases, followed by 5 deleted bases (gaps in the read relative to the reference), then 2 matched bases and finally 1 inserted base in the read compared to the reference.

When the letter ‘M’ is used in the CIGAR string to indicate alignment matches, without distinguishing between matches and mismatches (as opposed to using the extended CIGAR representation with ‘=’ for matches and ‘X’ for mismatches), the CIGAR string cannot be directly used to identify single-nucleotide polymorphisms (SNPs). This is because the ‘M’ operation represents both matches and mismatches to the reference sequence. For example, if the read sequence is ‘AATT’, the reference sequence is ‘AAAT’ and the alignment is represented as ‘4M’, the SNP at the third position is incorrectly treated as a match rather than a mismatch. Since BWA-MEM2 uses the ‘M’ notation in its alignments, the CIGAR string can only be directly used to calculate the number of insertions and deletions (indels) in each alignment, but not for identifying SNPs.

BWA-MEM2 includes an optional MD field in the SAM file that encodes information about mismatches between the read and the reference sequence. This field complements the CIGAR string by representing matches as integer values (without using the letters ‘M’, ‘X’, or ‘=’), mismatches as nucleotide letters and deletions as nucleotide letters preceded by the caret symbol (‘^’). The MD field does not provide information about insertions. For example, the MD string ‘5T2^TT10’ means:

- the first 5 bases perfectly match the reference,
- followed by a mismatch represented by the nucleotide ‘T’,
- then 2 perfect matches,
- followed by 2 deletions indicated by ‘^TT’,
- and ending with 10 perfect matches.

The number of SNPs in the alignment can be extracted by first calculating the number nucleotides in the MD string and thereafter subtracting this sum with the number of deletes in the CIGAR string. For example, if the CIGAR string is ‘8M2D3M2I7M’ and the equivalent MD string is ‘5T2^TT10’, the number of SNPs will be the number of nucleotides in the MD string, in total 3, subtracted with number of deletes in the CIGAR, in total 2, which gives 3-2 = 1 SNP.

### Variant analysis

The number of insertions, deletions and SNPs give an overview of the amount of variation that exists in the reads compared to the reference and could reflect the quality and amount of noise in the samples. Using the case study, these types of variations were extracted for the properly paired reads in each sample with separate analyses for Read 1 and Read 2 (Figure 6). The histograms illustrate the percentage of reads containing varying numbers of each variant type.

**Figure 6.**
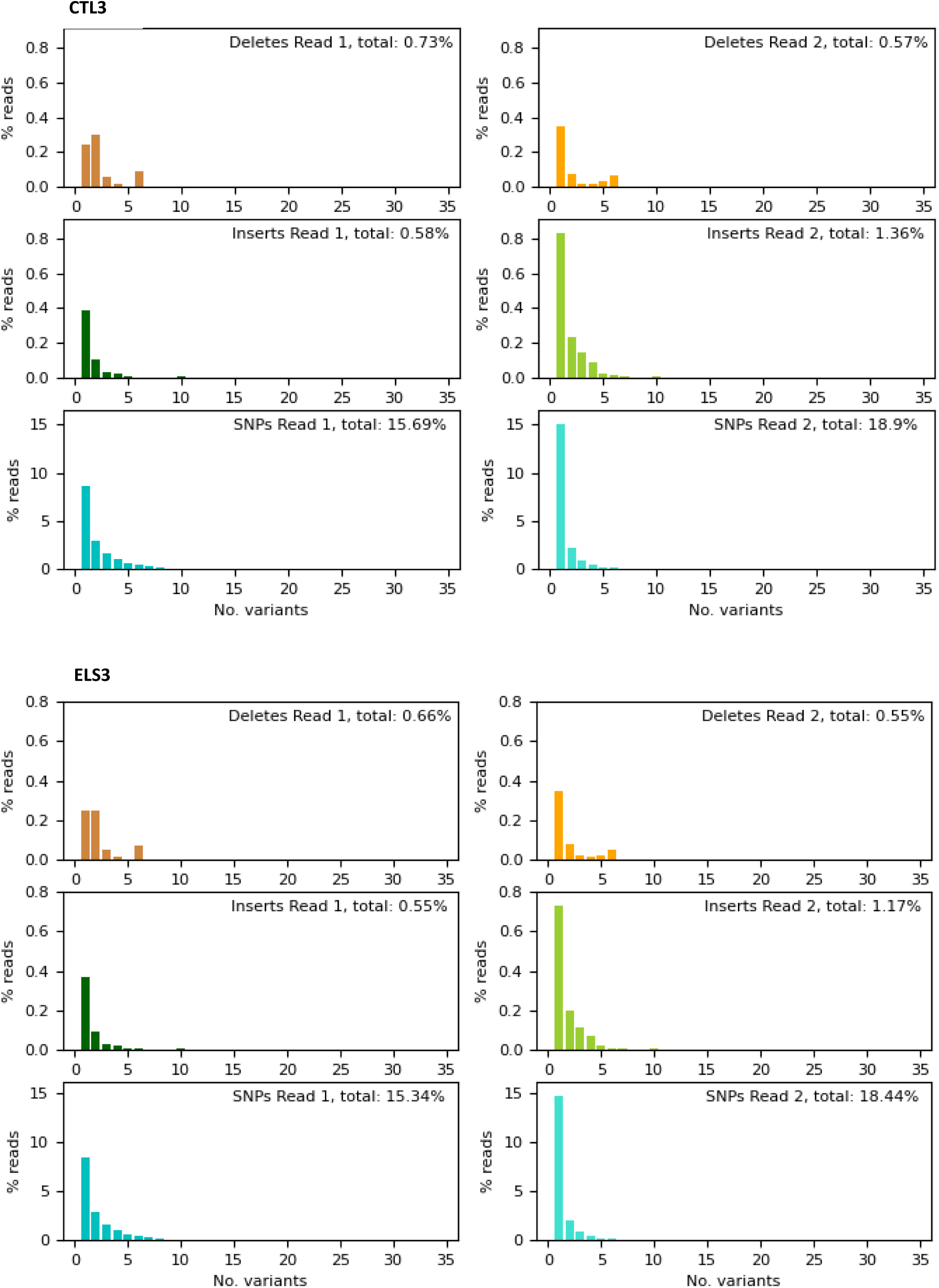
Histogram plots showing the amount of deletions, insertions and SNPs in the samples CTL3 (top) and ELS3 (bottom). The figures on left side for each sample depict results for first read population and on right side for second read population. The first two histograms for each sample depict deletions (brown-orange staples), second two histograms depict insertions (green-light green staples) and last two histograms depict SNPs (cyan-turquoise staples). The x-axis refers to the number of variations in the read and y-axis refers to the proportion (%) of reads having that many variations.

Deletions are relatively rare in both reads, with total frequencies ranging from approximately 0.55% to 0.73%. Most reads contain fewer than five deletions, and the percentage of reads with higher deletion counts declines sharply. Read 1 consistently shows a slightly higher deletion frequency than Read 2, suggesting minor differences in error profiles between the paired reads.

Insertions occur at a slightly higher rate in Read 2 compared to Read 1, with total insertion rates between 0.55% and 1.36%. The majority of reads have zero to five insertions, with a rapid decrease in frequency as the number of insertions increases. This pattern reflects typical sequencing characteristics where the reverse reads may have lower quality, leading to more insertions being detected.

On the other hand, SNPs are the most frequent variant type observed, present in approximately 15% to 19% of reads. Most reads carry between zero and five SNPs, with a peak at lower variant counts, and a few have >5 SNPs. To note, the number of variations is slightly higher in the second read population, most likely reflecting the fact that the sequence quality is generally lower for the second read due to the cumulative effects of sequencing chemistry limitations, library preparation artifacts, instrumental and environmental factors and the inherent challenges of base calling algorithms.

The similarity between the two figures demonstrates the reproducibility and consistency of variant detection across samples. The low levels of insertions and deletions reflect high-quality sequence alignments, as these variants often indicate sequencing or mapping errors. The higher prevalence of SNPs is expected and may represent genuine biological variation, RNA editing or technical noise. These results confirm the robustness of the BWA-MEM2 alignment algorithm and suggest that the data is suitable for further analyses, such as variant calling and expression profiling.

### Mapped Identity and Matched Identity

BWA-MEM2 includes the optional Alignment Score (AS) field in the SAM file, which quantifies the quality of a read’s alignment to the reference genome. This score can be used for assessing the quality and reliability of the alignment, with higher scores typically indicating better alignments. The AS field can also be used for distinguishing between multiple possible alignments for a read, as the highest score refers to the best alignment among several candidate alignments for the read.

The calculation of the AS depends on the specific alignment tool used, but generally incorporates factors such as the number of matches, mismatches and penalties for gaps (insertions and deletions), and sometimes also considering other factors such as base quality. For BWA-MEM2, the scoring system is defined by the following parameters: a match score of +1, a mismatch penalty of -4, a gap open penalty of -6 (applied to both insertions and deletions), and a gap extension penalty of -1 per base (also for both insertions and deletions). The AS is computed by adding the match score for each base that matches the reference, subtracting the mismatch penalty for each differing base, subtracting the gap open penalty whenever a gap is introduced and further subtracting the gap extension penalty for each base that extends the gap.

Chon and Huang (2021) have developed several concept-based mapping metrics to provide additional insights into alignment quality. These metrics can be useful for filtering out reads that do not align well to the reference sequence. One such metric is Mapped Identity, defined as the maximum summarized AS for a given read pair divided by the maximum possible summarized AS for the entire data set. For example, if the sequenced read length is 50 bases, the maximum AS for a single read corresponds to 50 (representing a perfect match of ‘50M’ in the CIGAR string). In paired-end sequencing, the maximum possible summarized AS for a read pair would hence be 100 (50 + 50) regarding this example. Suppose the first and second read receives an AS of 20 respectively 30, then the Mapped Identity is calculated as:

- Two paired-end reads are aligned and

- Read 1 has an AS of 20
- Read 2 has an AS of 30
- Maximum possible AS of 100 (50 + 50)

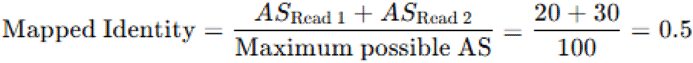

This metric provides an intuitive sense of how well the read pair aligns to the reference genome. As described by Chon and Huang (2021), “Conceptually, it is how well the pair of reads mapped anywhere to the provided reference.”

Another metric, introduced in this study, is Matched Identity, which reflects the proportion of perfectly matched bases in an alignment. The metric is calculated as the average of each AS divided by the number of matched bases (i.e., the number of ’M’s in the CIGAR string) for each read in a given read pair. Specifically, the AS for each read is divided by the corresponding number of matches and then the average of these two values is taken to represent the Matched Identity of the pair:

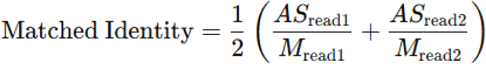

Using the same example as previously and assume the number of matches (’M’s) in the CIGAR string is 20 for the first read and 45 for the second read, then the Matched Identity is calculated as:

- Two paired-end reads are aligned and

Read 1 has an AS of 20 and number of M’s is 20
Read 2 has an AS of 30 and number of M’s is 45

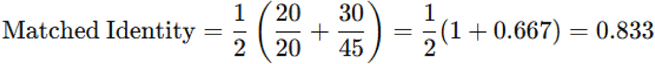

This metric quantifies how accurately the aligned portion of each read matches the reference sequence, by normalizing each read’s alignment score to the number of exact matches and averaging the two.

Mapped Identity is useful for filtering poorly aligned read pairs regardless of mismatch positions, while Matched Identity focuses on the alignment precision at the base level. Together, they provide complementary perspectives, as Mapped Identity gives on overall alignment success and Matched Identity gives a base-level accuracy. Applying thresholds to both metrics can help remove low-confidence alignments for more reliable downstream analyses.

### Metrics analysis

To make informed decisions about which thresholds to apply, binned histograms for each metric can be helpful (Figure 7). Each histogram bin shows the proportion of read pairs with scores either above or below a specified threshold. By selecting thresholds that range from very stringent to more lenient, the histograms illustrate the proportion of read pairs that would be retained or discarded at each level.

**Figure 7.**
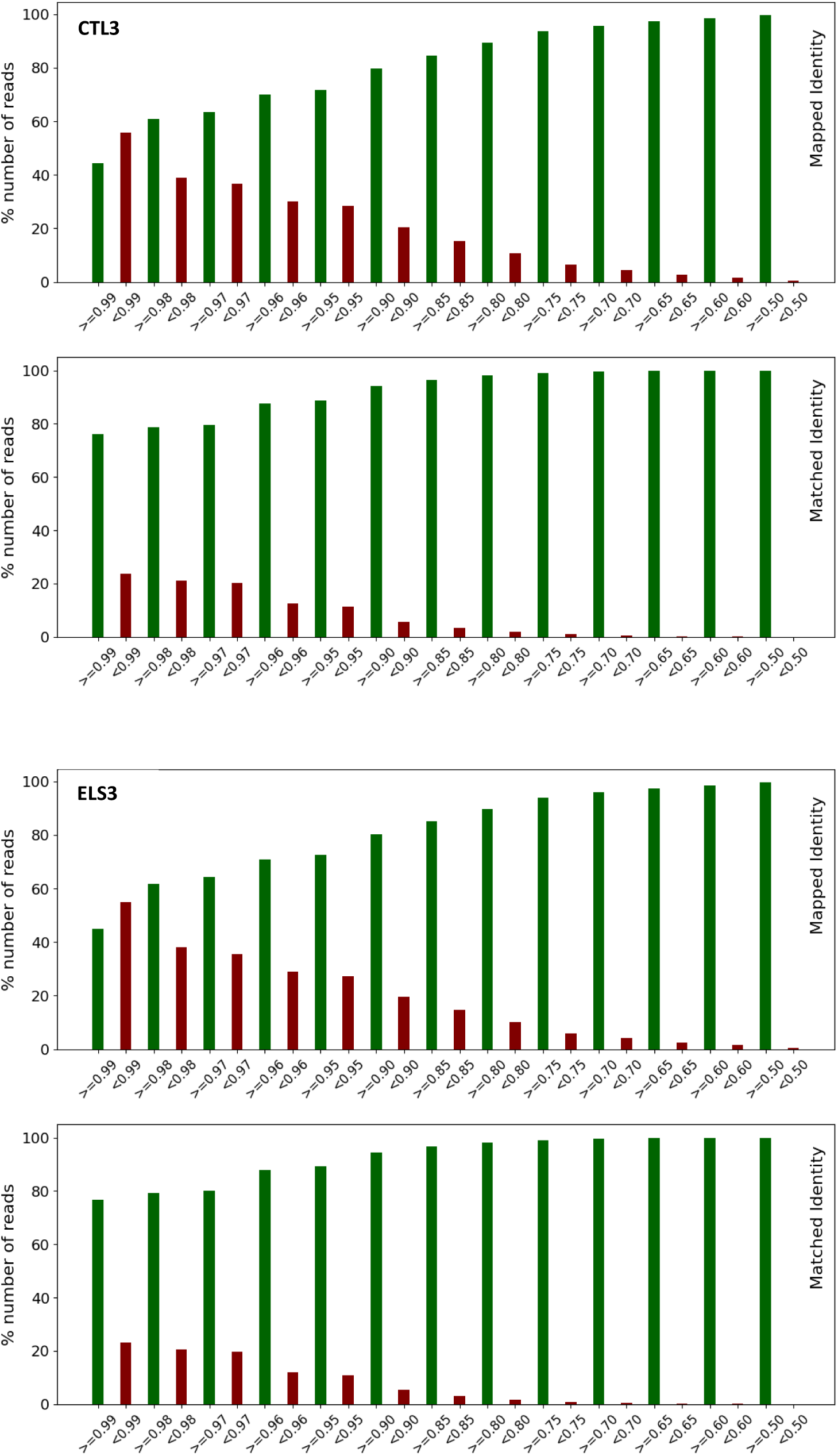
Binned histogram for each of the metrics, Mapped Identity respectively Matched Identity, for two of the samples, CTL3 respectively ELS3. The x-axis depicts different thresholds and the y-axis the % number of read pairs for which the threshold is satisfied.

In the dataset used as a case study, the histograms indicate that Mapped Identity is a stricter metric than Matched Identity, i.e., it discards more read pairs at the same threshold (Figure 7). For instance, at a threshold of ≥0.65, all read pairs are retained based on Matched Identity, while Mapped Identity requires a lower threshold of ≥0.4 to retain all. At a very stringent threshold of ≥0.99, about 20% of read pairs are discarded by Matched Identity, compared to around 60% by Mapped Identity.

The objective of applying these metrics is to discard read pairs with poor alignments, ensuring that only reliably mapped reads are retained for downstream analyses. Including too many read pairs with questionable mappings can obscure true biological signals and compromise the accuracy of the results. It is important to note that the *selection of threshold values ultimately lies with the user*, who must balance the trade-off between removing poorly aligned reads (to improve reliability) and retaining as many reads as possible (to preserve analytical depth and sensitivity). Below are alignment results from the mapping of reads in the CTL3 sample, provided to illustrate the metrics (Table 2). Maximum AS score is 76 for this data.

**Table 2.**
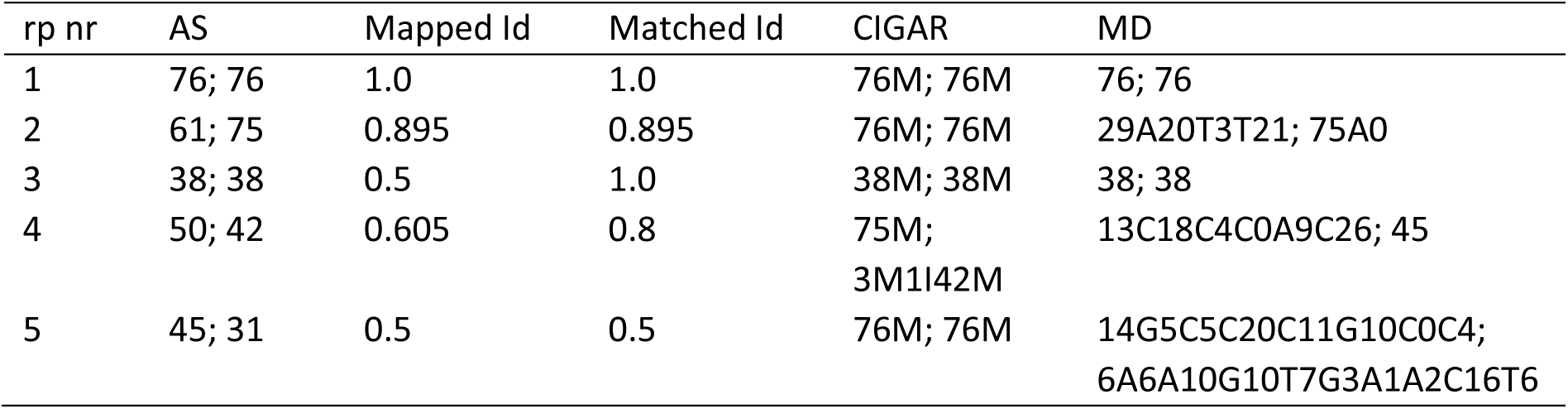
Examples of alignment results for sample CTL3. The columns give following information: rp nr, read pair number; AS, alignment score read 1 resp. read 2; Mapped Id, Mapped Identity score; Matched Id, Matched Identity score; CIGAR, CIGAR strings for read 1 resp. read 2; MD, MD strings for read 1 resp. read 2. Maximum AS score is 76.

| rp nr | AS | Mapped Id | Matched Id | CIGAR | MD |
| --- | --- | --- | --- | --- | --- |
| 1 | 76; 76 | 1.0 | 1.0 | 76M; 76M | 76; 76 |
| 2 | 61; 75 | 0.895 | 0.895 | 76M; 76M | 29A20T3T21; 75A0 |
| 3 | 38; 38 | 0.5 | 1.0 | 38M; 38M | 38; 38 |
| 4 | 50; 42 | 0.605 | 0.8 | 75M;<br>3M1I42M | 13C18C4C0A9C26; 45 |
| 5 | 45; 31 | 0.5 | 0.5 | 76M; 76M | 14G5C5C20C11G10C0C4;<br>6A6A10G10T7G3A1A2C16T6 |

For the first read pair (rp nr 1), both reads align perfectly, i.e., the CIGAR string is ‘76M’, yielding Mapped Identity and Matched Identity scores of 1.0. The AS reach the maximum value of 76 for both reads, and the CIGAR and MD strings show no variants relative to the reference sequence.

For the second read pair (rp nr 2), several variants are present relative to the reference sequence, as indicated by the MD strings. However, the CIGAR strings show no indels in the alignments, only matches. Since the number of matches in the CIGAR string equals the maximum AS, the metrics will yield the same score. While a few variants (three in the first read and one in the second) slightly reduce the metric scores, they are not numerous enough to warrant removing the alignment from the results.

For the third read pair (rp nr 3), the Matched Identity gives a perfect score, as the alignment consists of only matches and there are no variants indicated. However, the length of the matched sequences is relatively short, only 38 bases, and therefore the Mapped Identity score is low. Hence, in this case, the Mapped Identity score reflects a short mapping length, but otherwise a perfect alignment.

For the fourth read pair (rp nr 4), the CIGAR string reveals several SNPs relative to the reference sequence in the first read, but with a substantial alignment length. The second read, however, shows a shorter alignment length and an insertion, though it still retains a good number of matches (M’s). These factors contribute to a more significant reduction in the Mapped Identity score compared to the Matched Identity score. This read pair exemplifies the impact of short alignment length combined with the presence of multiple variants.

The fifth read pair (rp nr 5), exhibits significant variations relative to the reference sequence, despite both reads having maximum alignment lengths. Both reads contain multiple SNPs, which reduces both the Mapped Identity score and the Matched Identity score. This result is less trustable and should preferably be removed before subsequent analyses.

### Metrics based filtering

Different thresholds need to be set for the two metrics, as indicated by the binned histograms. Our aim is to remove only read pairs with very low scores, thereby filtering out dubious mappings while retaining reliable ones. Based on this, the thresholds were set to 0.6 for Mapped Identity and 0.8 for Matched Identity. However, it is recommended to test various thresholds and assess their impact on downstream analyses, such as the number of identified differentially expressed genes. This filtering step results in the removal of an additional ∼2–3% of reads from the samples (Table 3). The low percentage of reads removed reflects the overall high quality of the sequencing data and the alignments. Most reads passed earlier quality control steps and aligned well to the reference genome, therefore only a small fraction fell below the set thresholds. Additionally, the thresholds were set to remove only the most dubious alignments, ensuring that the majority of trustworthy reads are retained for reliable downstream analyses.

**Table 3.** Statistics regarding: Total, total number of reads sequenced in each sample (counting reads sequenced from both ends); Properly paired, proportion of reads assigned the SAM flag 99, 83, 163 or 147; Other, proportion of reads assigned any other SAM flag value; Properly paired, quality alignm., reads in an alignment satisfying set thresholds for Mapped Identity and Matched Identity.

|  | CTL1 | CTL2 | CTL3 | ELS1 | ELS2 | ELS3 |
| --- | --- | --- | --- | --- | --- | --- |
| Total | 46,894,541 | 43,859,473 | 44,733,495 | 50,107,279 | 54,711,209 | 49,670,950 |
| Properly paired | 78.5% | 78.9% | 78.9% | 78.6% | 79.0% | 79.0% |
| Other | 21.5% | 21.1% | 21.1% | 21.4% | 21.0% | 21.0% |
| Properly paired,<br>quality alignm. | 76.4% | 76.7% | 76.5% | 76.2% | 76.7% | 76.8% |

To provide deeper insight into the two metrics additional scatterplots were generated for each sample. These plots display the relationship between each metric’s score and the number of variants in each read pair. Specifically, the total number of deletions, insertions, SNPs and overall variants were summed for each read pair. and then plotted against the Mapped Identity and Matched Identity scores (Figure 8). The filtering process has a significant impact on SNPs, with reads containing more than approximately 15 SNPs being discarded. In contrast, deletions and insertions are less affected, with only some reads containing these variants being filtered out.

**Figure 8.**
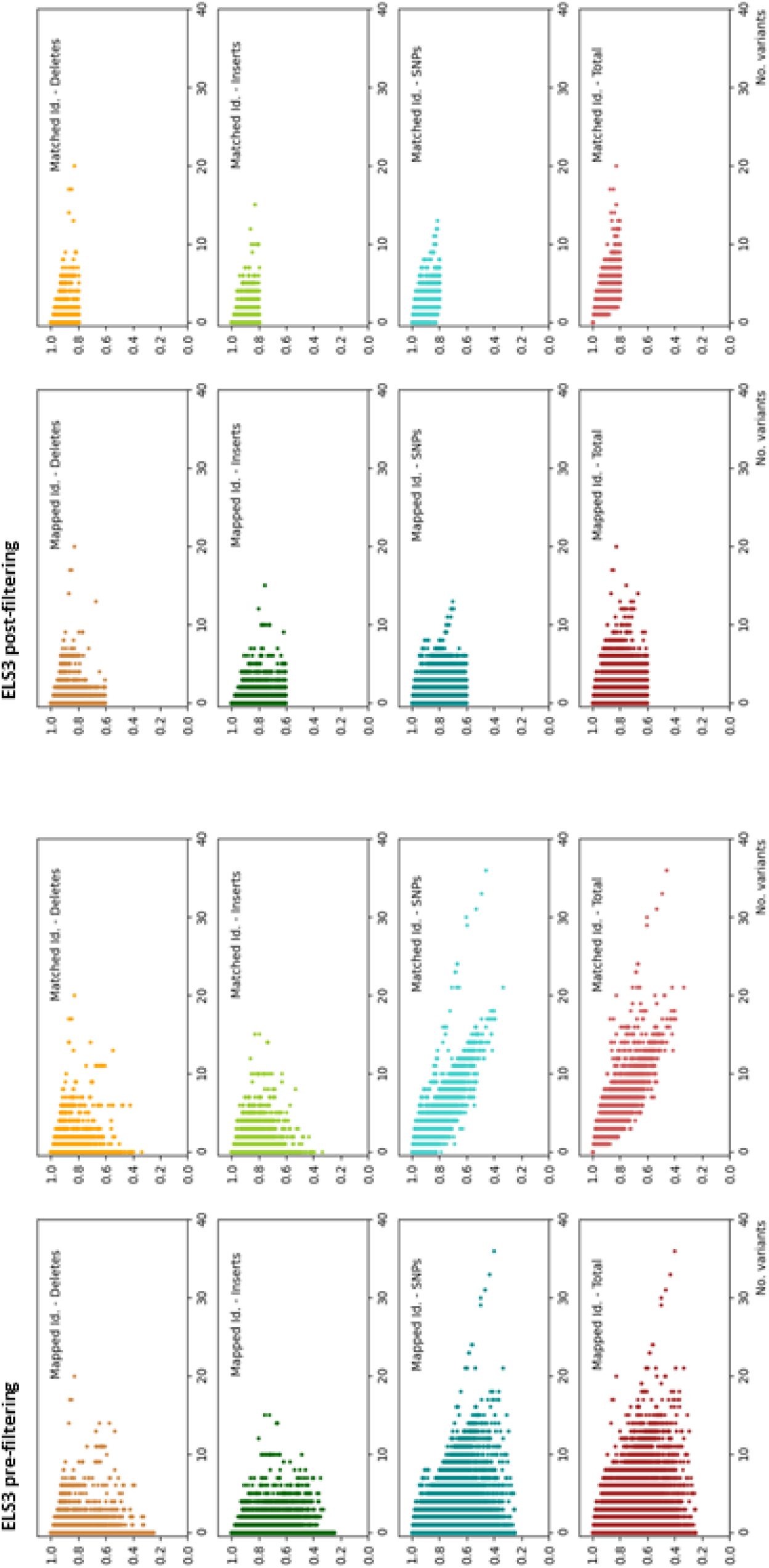
Scatterplots generated for sample ELS3 pre- and post-filtering, left vs. right hand figure, using thresholds Mapped Identity >= 0.6 and Matched Identity >= 0.8. These plots display the relationship between each metric’s score and the number of variants in each read pair, from top to bottom: deletes, inserts, SNPs and overall. The x-axis depicts number of variants in both reads for a read pair and y-axis depicts the metric score.

### Chromosomal distribution

By analyzing the distribution of mapped reads per chromosome, we can gain insights into the quality of the RNA-seq data, biological characteristics of the sample and potential technical issues that may need to be addressed. For instance, read counts per chromosome can reflect gene density, where chromosomes with more actively transcribed genes tend to have higher read coverage. Additionally, certain chromosomes may be more transcriptionally active in specific tissues or cell types, offering biologically relevant information. A high proportion of reads mapping to the mitochondrial genome may suggest elevated metabolic activity or could signal sample quality concerns, such as cytoplasmic RNA degradation, which leaves more stable mitochondrial RNA intact. Differences in chromosomal read distribution between samples may indicate biases introduced during RNA extraction, library preparation, or sequencing, as well as potential alignment problems like incorrect mapping by the aligner or challenges in mapping reads from homologous regions. Furthermore, reads mapping to unplaced or unlocalized sequences can help identify novel transcripts or structural variations, adding another layer of biological insight.

Figure 9 presents histograms of read pairs per chromosome for samples CTL3 and ELS3. The distribution of mapped read pairs per chromosome shows that most reads originate from the mitochondrial genome, followed by chromosomes 1, 2, 7, and 11, which have similar read counts. The remaining chromosomes display a gradual decrease in read numbers, with relatively few reads mapping to chromosomes X and Y. Additionally, a small number of reads map to unplaced and unlocalized sequences. All samples exhibit a similar distribution pattern, indicating no issues during sample preparation, sequencing, or alignment. Furthermore, comparison between CTL and ELS samples reveals no significant differences in chromosomal read distribution, suggesting that early life stress does not specifically affect the representation of reads from any particular chromosome.

**Figure 9.**
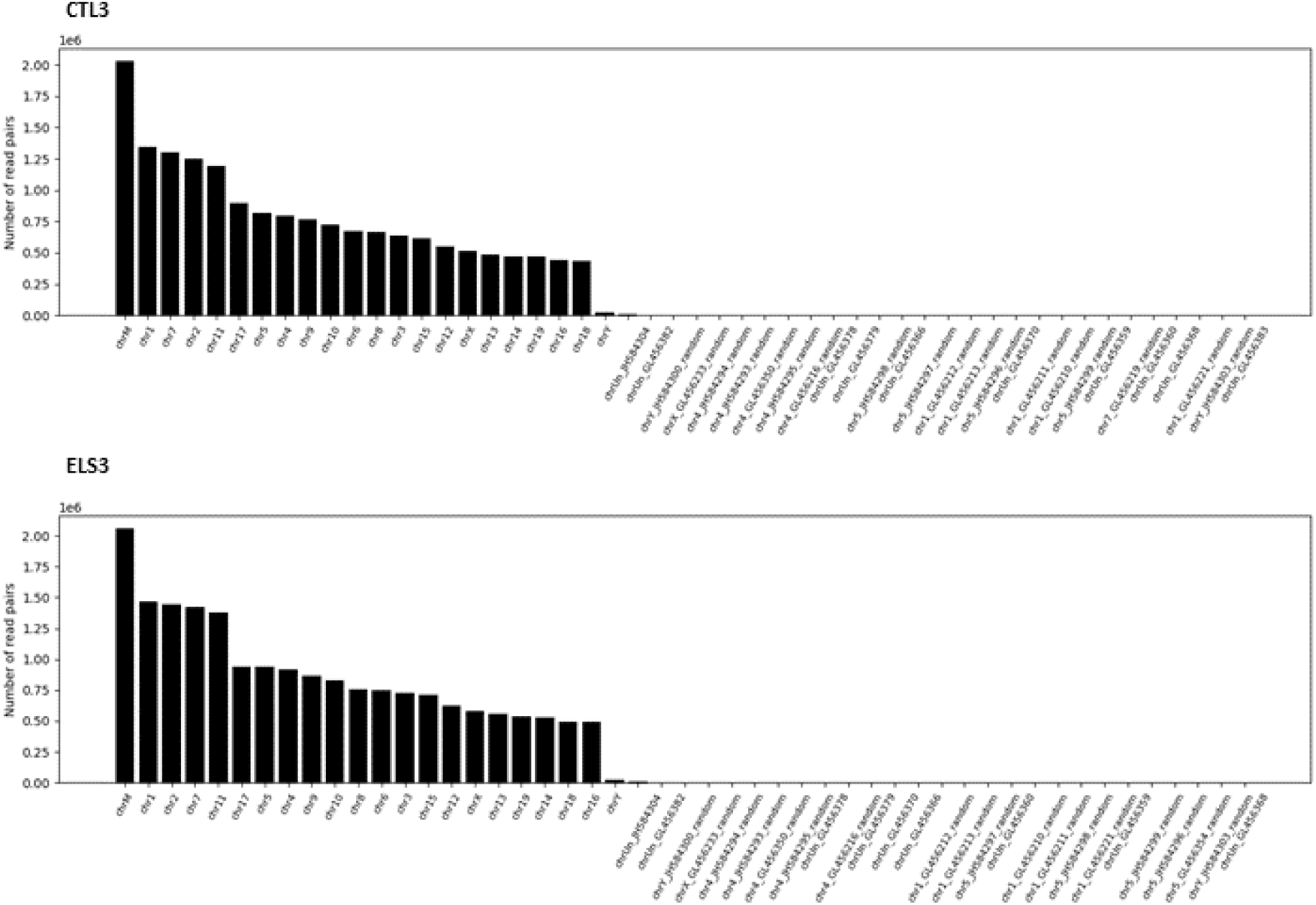
Histograms displaying number of mapped read pairs per chromosome, unplaced and unlocalized sequences in the reference. Sample CTL3 top and ELS3 bottom histogram, respectively. The x-axis depicts chromosome or sequence number and y-axis number of read pairs.

### Counting chromosomal mappings

To gain a more detailed understanding of read alignments across chromosomes, we can count the number of reads mapped to each base pair (bp) position using information from the SAM file. For example, the hypothetical alignment data in Table 4 illustrates how reads are mapped, showing both the starting position of each alignment and the alignment pattern as described by the CIGAR string. The hypothetical alignment shows results for three reads. Read r1 starts at position 53 and aligns perfectly for 6 bases (6M). Read r2 starts at position 52 and includes a deletion after 2 matched bases (2M1D3M). Read r3 also starts at position 52 and contains an insertion after 4 matched bases (4M1I2M). These examples illustrate different types of alignment operations represented by the CIGAR strings.

**Table 4.** Hypothetical alignment results. Read nr, fictive read number ID; Start pos, chromosomal start positions for alignment; CIGAR, hypothetical CIGAR string for alignment.

| Read nr | Start pos | CIGAR |
| --- | --- | --- |
| r1 | 53 | 6M |
| r2 | 52 | 2M1D3M |
| r3 | 52 | 4M1I2M |

In this fictive example, for the first read the CIGAR gives six matches (6M) starting at chromosomal position 53 and, therefore, therefore each position from 53 to 58 we can be counted as covered once, +1, (Table 5).

**Table 5.** Counts from read alignments. Chr pos, chromosomal position; Read nr, fictive read number ID; Counts, aggregated counts for each chromosomal position.

| Chr pos |  |  |  |  |  |  |  |  |
| --- | --- | --- | --- | --- | --- | --- | --- | --- |
| Read nr | 52 | 53 | 54 | 55 | 55.1 | 56 | 57 | 58 |
| r1 |  | 1 | 1 | 1 |  | 1 | 1 | 1 |
| r2 | 1 | 1 |  | 1 |  | 1 | 1 |  |
| r3 | 1 | 1 | 1 | 1 | 1 | 1 | 1 |  |
| Counts | 2 | 3 | 2 | 3 | 1 | 3 | 3 | 1 |

For the second read, the CIGAR contains two matches, followed by a deletion and then three more matches. For this read, we count +1 for positions 52 and 53. However, due to the deletion at position 54, no count is recorded there. Counting resumes with +1 for positions 55 to 57, corresponding to the three matches after the deletion.

For the third read, there are four matches, followed by an insertion not present in the other reads, and then two more matches. We count +1 for the initial four matches spanning positions 52 to 55. Because of the insertion after position 55, we introduce an additional position, labeled 55.1, to represent this inserted base relative to the reference genome. The other reads do not contribute to position 55.1, as they have no insertion there. Finally, we count +1 for positions 56 and 57, which correspond to the two matches following the insertion.

In summary, we aggregate the counts at each chromosomal position to determine how many reads are mapped to that specific location. In this example, the distribution of mapped reads varies across positions, with a peak count of 3 observed at positions 53, 55, 56, and 57. The lowest counts occur at position 55.1, corresponding to the insertion site, and position 58. This variation reflects the differences in how individual reads align, including the presence of insertions or deletions that shift read coverage at certain bases. Such detailed positional information is important for identifying regions of high or low coverage, which can inform downstream analyses like variant detection or expression quantification. Furthermore, accurately accounting for insertions by introducing fractional positions (like 55.1) helps maintain the integrity of mapping data and ensures that all bases, including those not present in the reference, are properly represented in the coverage profile (Table 5).

### mtDNA mappings

Using the approach described above, we can examine the mapping results for mitochondrial DNA (mtDNA) across all samples by plotting them together in a single figure (Figure 10). The mitochondrial genome spans 16,500 base pairs and contains 37 genes, all tightly packed without introns, meaning that each gene starts immediately after the previous one ends. The figure shows coverage plots for each sample, where the x-axis represents genomic positions and the y-axis shows the number of reads mapped to each position. Gene start positions are indicated with green horizontal lines whereas gene end positions are indicated with red horizontal lines. However, because the genes are closely packed together, the red line marking the end of one gene often overlaps with the green line marking the start of the next gene, making them hard to distinguish visually.

**Figure 10.**
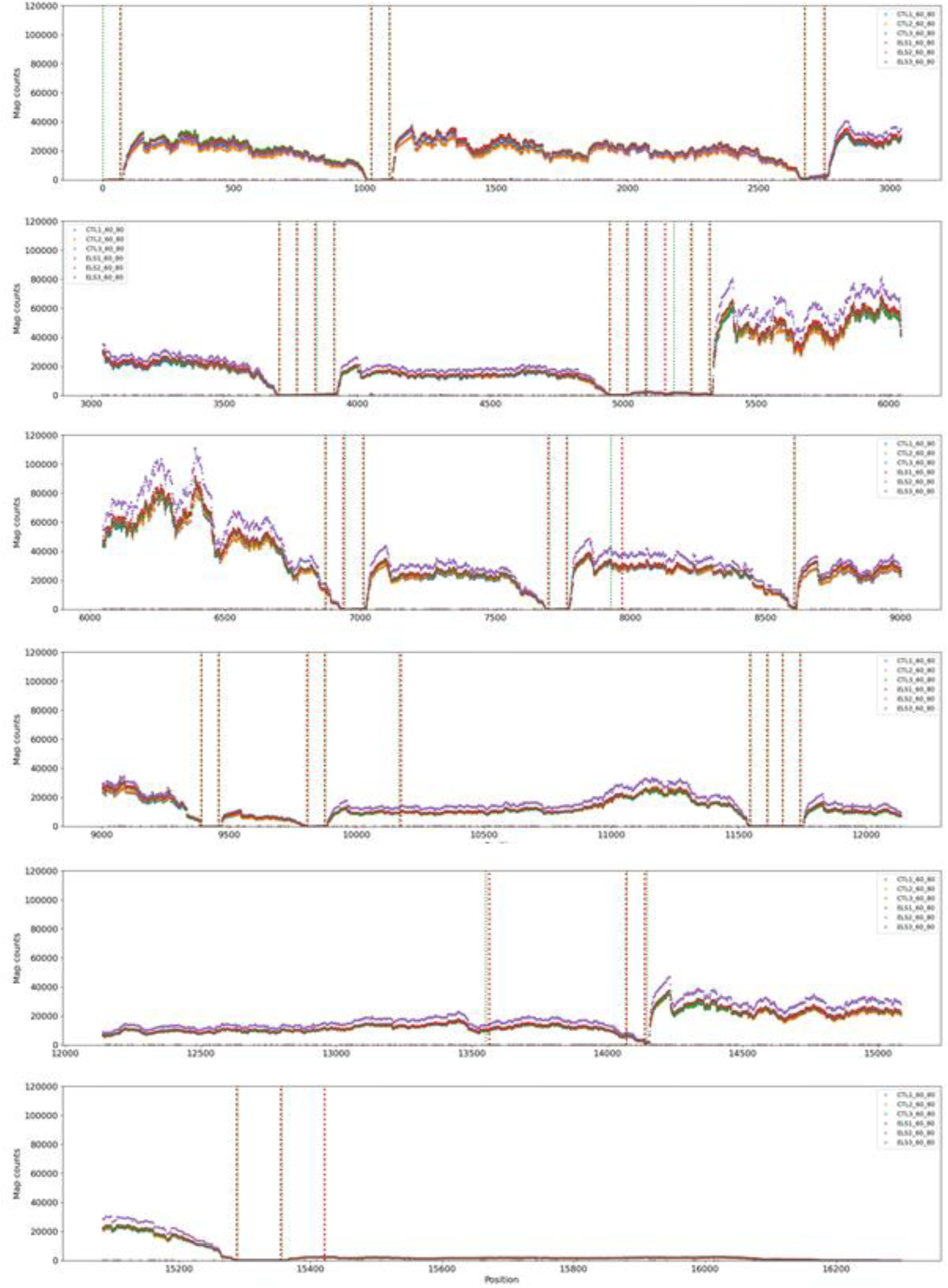
Each curve represents the read mapping coverage of one sample across the 16,500 base pairs (bp) of the mitochondrial DNA (mtDNA). The x-axis shows the genomic position, while the y-axis indicates the number of reads mapped to each position. Gene start positions are marked by green horizontal lines and gene end positions by red horizontal lines. Due to the compact nature of the mitochondrial genome, with genes located immediately next to each other, start and end markers may overlap and appear visually indistinct.

Across all samples, 2,525 insertion sites were detected throughout the mtDNA. Most insertions occur rarely: 56% of positions have only one insertion, while about 1% have more than 90 insertions. Similarly, 2,672 deletions were identified, with 52% having a single occurrence and only 0.3% showing more than 90. No specific position or region stands out with unusually high insertion or deletion activity, indicating a generally even distribution of such variants across the mitochondrial genome.

The read mapping coverage appears to be fairly consistent among the different samples, as indicated by the overlapping lines, and similar trends in coverage are observed across these samples. This consistency is further validated by calculating the Kendall correlation between each treated sample and control sample (Table 6). The lowest correlation observed between any treated and control sample is 0.91, indicating a strong correlation in mapping coverage across the entire genome.

**Table 6.** Calculated Kendall correlation between each treated and control sample for mtDNA. Treated, treated sample; Control, control sample; Kendall, calculated Kendall correlation between the two samples.

| Treated | Control | Kendall |
| --- | --- | --- |
| ELS1 | CTL1 | 0.96 |
| ELS1 | CTL2 | 0.95 |
| ELS1 | CTL3 | 0.97 |
| ELS2 | CTL1 | 0.93 |
| ELS2 | CTL2 | 0.94 |
| ELS2 | CTL3 | 0.91 |
| ELS3 | CTL1 | 0.96 |
| ELS3 | CTL2 | 0.94 |
| ELS3 | CTL3 | 0.97 |

There are noticeable drops in read coverage at certain positions in the mtDNA, which corresponds to deletions and insertions in individual samples (Figure 10). While the coverage is mostly consistent, some regions show slight variations in read counts between the samples. This could be due to differences in how the RNA from these regions was amplified, sequenced, aligned or even differences in gene expression. For some genes, no reads have been mapped, indicating that they may either be unexpressed, have been missed during sequencing or been mis-aligned by the aligner. There is also natural variation in gene expression levels, where some genes are lowly expressed, i.e., having a generally low number of counts for all positions, whereas some genes are highly expressed with a large number of mapped reads. For example, the gene COX1 on positions 5328 to 6972 shows highest expression and its protein is a key enzyme in aerobic metabolism.

### nDNA mappings

Chromosomal nuclear DNA (nDNA), on the other hand, contains both exons and introns. During transcription, only the exons contribute to mRNA synthesis, while the introns are spliced out and thus not represented in RNA sequencing data. As a result, mapping RNA-Seq reads to nuclear chromosomes produces regions with and without read alignments. The next step in NGS analysis is to extract the regions with aligned reads on each chromosome and analyze them in greater detail.

Figure 11 shows the mappings for three regions on chromosome 1: positions 3003336-3003558, 3199741-3207767 and 9600952-9602461, respectively. The number of aligned reads for the first region is extremely low, with only samples CTL1 and CTL2 having one mapped read each. The second region, on the other hand, has more mapped reads across all samples, with at least one sample having at least one position with more than 50 mapped reads. In this region, sample ELS2 has the highest number of mapped reads at a single position; 56 reads at both positions 3,200,296 and 3,200,297.

**Figure 11.**
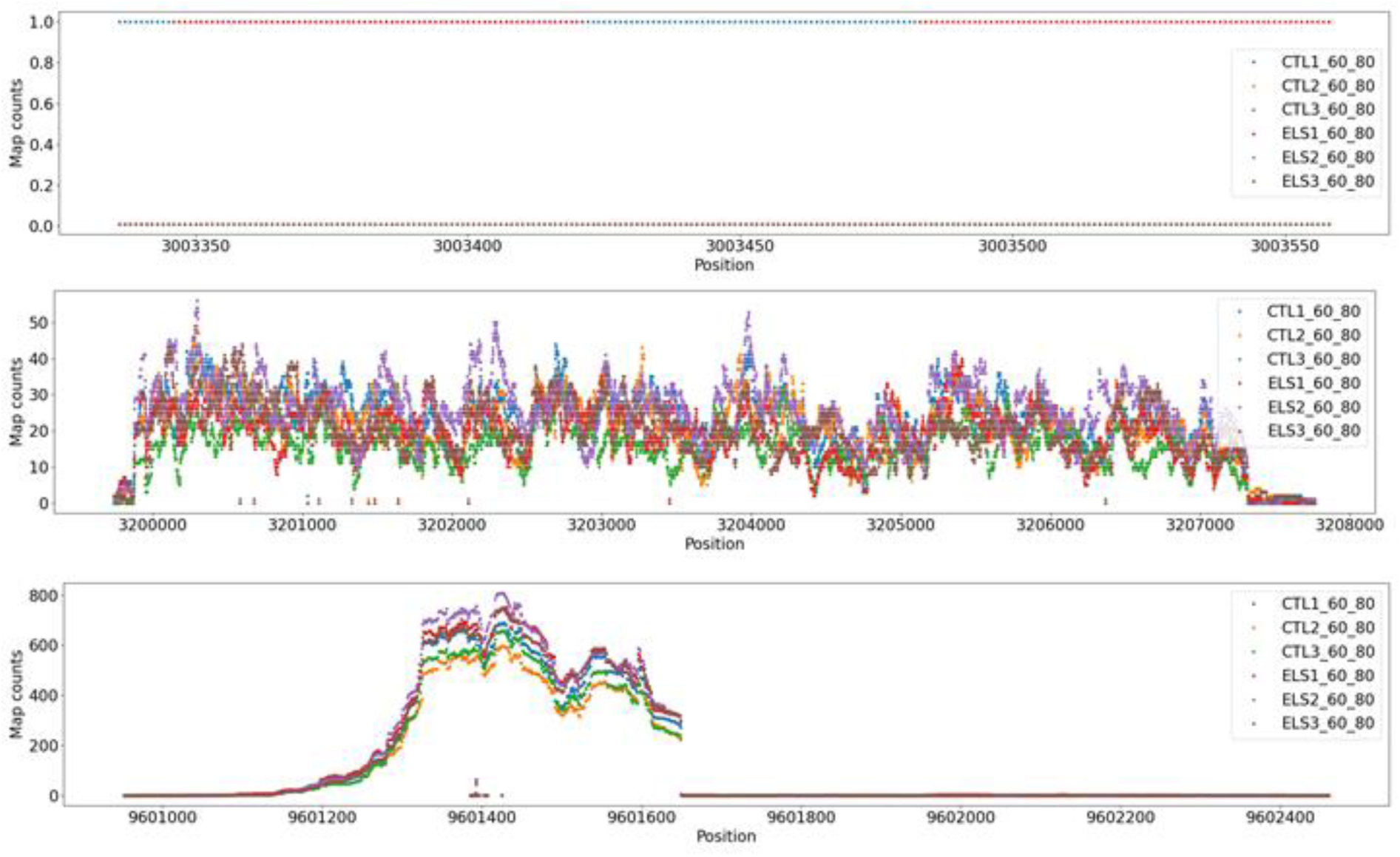
Read counts for four regions on chromosome 1. In order from top to bottom, positions 3003336-3003558, 3199741-3207767 and 9600952-9602461, respectively. The x-axis depict position in the region and y-axis number of counts.

The third region shows even more mapped reads, with the maximum mapped read count appears to be just above 800 and which occurs near the peak of the coverage region (∼position 9601400– 9601450). This peak is highest for the ELS2 sample, shown in purple, which rises slightly above the other samples at the peak. A prominent peak in read coverage is observed between positions ∼9601200 and ∼9601650 across all samples, indicating a region of high transcription or enrichment. Both CTL and ELS samples follow similar coverage patterns for this region, though subtle differences in peak height and distribution suggest potential variability between groups. Notably, there is a minor dip in coverage around position ∼9601425, consistently present across all samples. Outside the main peak region, particularly beyond position ∼9601700, read counts drop sharply and remain close to zero, indicating little to no mapping in this part. This pattern suggests that the region of interest is highly localized and may correspond to a functional genomic element such as an exon, promoter or regulatory sequence.

Figure 11 highlights that the region between positions 3199741 and 3207767 shows greater variability in expression levels across samples compared to mtDNA. This observation is supported by the Kendall correlation, which ranges from 0.39 to 0.52 between treated and control samples (Table 7). The low correlation between sample groups suggests a possible difference in expression levels under treatment versus normal conditions. However, the similarly low Kendall correlation (0.43-0.54) within each sample group indicates that the data does not provide strong evidence for significant deregulation of this region due to treatment (Table 8). On the contrary, significant deregulation would have been indicated by a strong correlation between samples within the treatment group (within-group similarity) and low correlation between samples from the two different groups (between-group similarity).

**Table 7.** Calculated Kendall correlation between each treated and control sample for position 3199741-3207767 on chromosome 1. Treated, treated sample; Control, control sample; Kendall, calculated Kendall correlation between the two samples.

| Treated | Control | Kendall |
| --- | --- | --- |
| ELS1 | CTL1 | 0.52 |
| ELS1 | CTL2 | 0.49 |
| ELS1 | CTL3 | 0.43 |
| ELS2 | CTL1 | 0.51 |
| ELS2 | CTL2 | 0.46 |
| ELS2 | CTL3 | 0.39 |
| ELS3 | CTL1 | 0.48 |
| ELS3 | CTL2 | 0.48 |
| ELS3 | CTL3 | 0.39 |

**Table 8.** Calculated Kendall correlation between two samples from the same group for position 3199741-3207767 on chromosome 1. Sample 1, first sample selected from group; Sample 2, second sample selected from same group as first sample; Kendall, calculated Kendall correlation between the two samples.

| Sample 1 | Sample 2 | Kendall |
| --- | --- | --- |
| ELS1 | ELS2 | 0.47 |
| ELS1 | ELS3 | 0.49 |
| ELS2 | ELS3 | 0.43 |
| CTL1 | CTL2 | 0.54 |
| CTL1 | CTL3 | 0.51 |
| CTL2 | CTL3 | 0.47 |

Regarding the region between positions 9600952 and 9602461, Kendall correlations show moderate to high relationships between the ELS and CTL sample groups as they range from 0.61 to 0.81 (Table 9). The highest correlation can be observed between ELS2 and CTL3 (0.814). In contrast, ELS3 and CTL2 (0.609) has the lowest cross-group correlation, indicating more divergence. Within the ELS group, correlations ranged from 0.66 to 0.75, with ELS2 and ELS3 showing the strongest correlation (0.749; Table 10). For this region, CTL samples have slightly generally higher within-group consistency, with correlations from 0.76 to 0.87 and they exhibit stronger internal coherence than ELS samples. The variability among ELS samples could possibly reflect greater expression heterogeneity in that group. Moreover, most ELS-CTL correlations fall in the 0.7-0.8 range, indicating moderate similarity across groups.

**Table 9.**
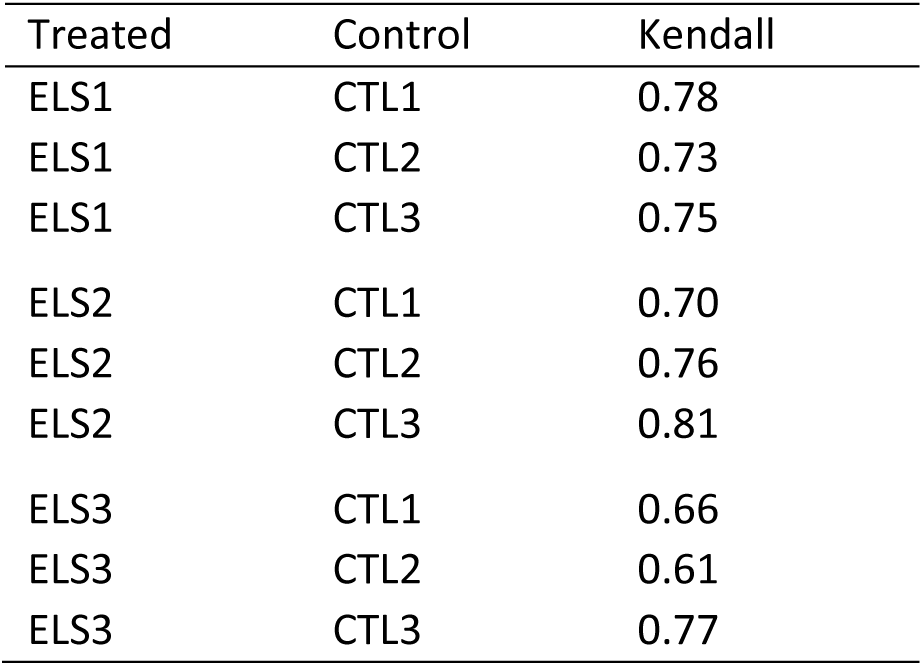
Calculated Kendall correlation between each treated and control sample for position 9600952-9602461 on chromosome 1. Treated, treated sample; Control, control sample; Kendall, calculated Kendall correlation between the two samples.

**Table 10.** Calculated Kendall correlation between two samples from the same group for position 3199741-3207767 on chromosome 1. Sample 1, first sample selected from group; Sample 2, second sample selected from same group as first sample; Kendall, calculated Kendall correlation between the two samples.

| Sample 1 | Sample 2 | Kendall |
| --- | --- | --- |
| ELS1 | ELS2 | 0.66 |
| ELS1 | ELS3 | 0.68 |
| ELS2 | ELS3 | 0.75 |
| CTL1 | CTL2 | 0.76 |
| CTL1 | CTL3 | 0.79 |
| CTL2 | CTL3 | 0.87 |

RNA-Seq mapping to nuclear DNA reveals uneven and diverse read distributions, largely due to the presence of both exons and introns and varying transcription levels. As illustrated here, one region displays minimal expression, another shows moderate expression with sample-to-sample variability, and a third exhibits a strong, localized transcriptional peak. Importantly, expression variability across genomic regions can differ markedly and not all observed differences are necessarily attributable to treatment effects. Some regions demonstrate consistent expression across conditions, while others are more variable, likely reflecting underlying biological complexity rather than direct treatment-induced changes.

### Count filtering

To improve data quality, mapped regions can be filtered based on read count levels, allowing lowly expressed regions to be excluded from further analysis. A common strategy is to retain only regions where at least one sample shows a minimum number of reads at any position, with the threshold set by the researcher. For example, applying a threshold of ≥10 counts at any position in a region and in at least one sample reduces the number of regions on chromosome 1 from 121,732 to 5,159. Raising the threshold to ≥20 counts further reduces this number to 3,327 (Figure 12).

**Figure 12.**
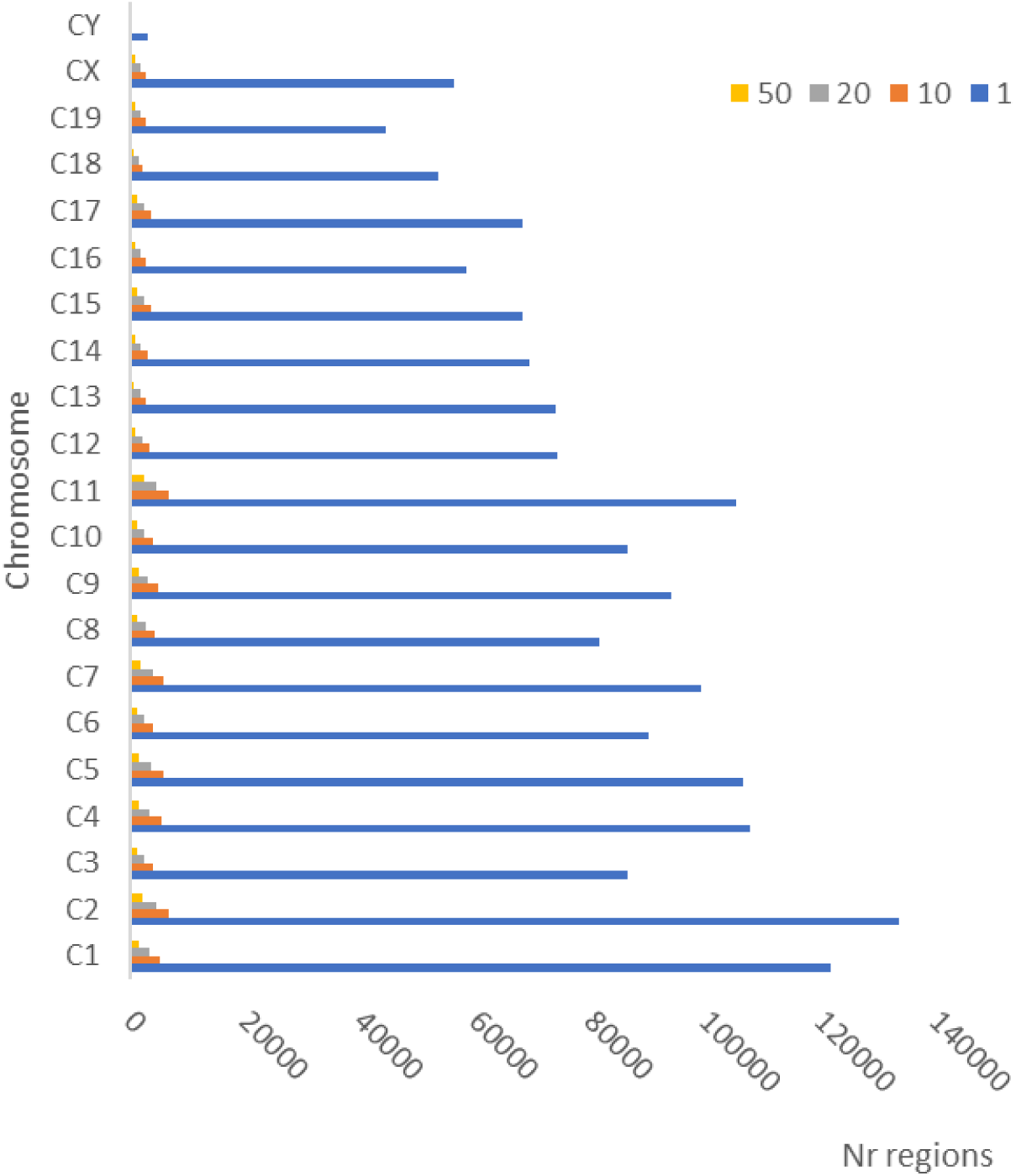
The number of derived regions for each chromosome depending on the selected count threshold. The x-axis represents the number of derived regions, while the y-axis corresponds to the different chromosomes. The colors indicate the chosen threshold for the minimum number of counts at any position within the region in at least one sample.

By setting user-defined thresholds for read counts, regions with low expression can be excluded, allowing researchers to focus on biologically relevant areas and retain only the most informative regions for downstream analysis. Figure 12 shows how the number of retained regions per chromosome varies with different count thresholds. On average, 95 ± 0.8% of regions across chromosomes have fewer than 10 counts at any position in any sample.

When setting a threshold, it is important to consider the biological context: a threshold that is too high may discard meaningful but moderately expressed regions, while a threshold that is too low may retain uninformative regions. Because expression levels can vary between samples or experimental conditions, the threshold should also account for this variability to avoid introducing bias. Examining the read count distribution beforehand can help inform an appropriate cutoff. Lastly, the sequencing depth, i.e., the total number of sequenced fragments, will influence the overall number of mapped reads and should be considered when selecting the threshold.

## Materials and Methods

### FastQC and MultiQC

FastQ files from the Gene Expression Omnibus dataset GSE180055 were loaded into Galaxy Europe server (https://usegalaxy.eu/) using the ‘Faster Download and Extract Reads in FASTQ format from NCBI SRA’. Quality control of the paired-end FASTQ files was performed using FastQC (Andrews, 2010), which generated individual reports for forward and reverse reads. MultiQC (Ewels et al., 2016) was thereafter used to aggregate these reports into a comprehensive summary for each sample. Quality assessments were based on the FastQC analyses: Per Base Sequence Quality, Per Sequence Quality Scores and Sequence Length Distribution. These metrics were evaluated for all samples, with detailed illustrations from two representative mouse samples, ELS3 and CTL3 (Usui et al., 2021).

### Read alignment

Read alignment with the FastQ files was performed using BWA-MEM2 (Li & Durbin, 2010) on the Galaxy Europe server using the available reference genome mm10 for *Mus musculus*. The tool was executed with default parameters, except for sorting alignments by read name instead of chromosomal position. Due to the high quality of the dataset and minimal adapter contamination observed in FastQC/MultiQC reports, no trimming was performed prior to alignment; instead, BWA-MEM2’s soft clipping feature was relied upon to remove low-quality or contaminant bases at read ends. Output alignments were generated in BAM format, with one file per sample containing both forward and reverse reads.

### BAM file parsing

To extract and analyze properly paired sequencing reads from the alignment data, a custom Python script was developed that utilizes the `bamnostic` library for parsing BAM files. During execution, the BAM file is read line by line and extracts properly paired reads, as defined by the SAM flags 83, 99, 147, and 163. Reads reported with other flags were not properly paired and these reads were discarded from subsequent analyses. For each properly paired read, the mapping quality, CIGAR string, reference positions, alignment score (AS tag) and mismatch information (MD tag) were extracted. These data were grouped by read identifiers and organized into read-pair batches. Processed alignment data were saved to an output file for downstream analysis. The script also generated summary statistics in form of total number of reads, number of properly paired reads, number of reads not properly paired and visualization of flag distribution for not properly paired reads.

### Variant analysis

Using a custom Python script and the processed alignment data generated in previous step, the number of insertions and deletions for each read alignment was directly extracted from the CIGAR strings. The number of SNPs in each read alignment was calculated based on the information given in the CIGAR and MD strings, according to:

- *N_MD_* = total number of nucleotide mismatches (substitutions) indicated in the MD string,
- *D_CIGAR_ =* total number of deleted bases indicated in the CIGAR string.

Then the number of Single Nucleotide Polymorphisms (SNPs) is:

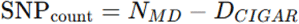

Histogram plots were thereafter generated for each sample showing the number of variations versus the proportion (%) of reads having that many variations.

### Mapped Identity and Matched Identity scores

To quantify alignment accuracy for paired-end reads, two scoring functions were implemented in a custom Python script. The scoring functions rely on the alignment score (AS) reported by BWA-MEM2 and, additionally, for Matched Identity the number of matches (M) in the alignments.

Mapped Identity is calculated as:

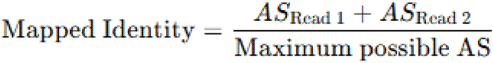

Matched Identity is calculated as:

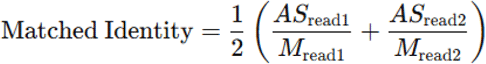

To assess read alignment quality, the distributions of the two were visualized by plotting their binned histograms. For each metric, reads were binned according to defined score thresholds:
≥0.99, <0.99, ≥0.98, <0.98, ..., ≥0.50, <0.50.

This resulted in 26 bins, with alternating bars representing the number of reads that either met (≥ threshold) or did not meet (< threshold) the specified cutoff. Bar heights correspond to the percentage of total reads falling within each bin, allowing for a visual quantification of score distribution across increasing stringency levels. The bars were color-coded with dark green bars indicating the proportion of reads satisfying each threshold (e.g., MI ≥ 0.99) and maroon bars represent reads that failed to meet the threshold (e.g., MI < 0.99).

### Metrics based filtering

Using a custom Python script, read pairs were subsequently filtered based on predefined thresholds of Mapped Identity ≥ 0.6 and Matched Identity ≥ 0.8, respectively. The number and percentage of read pairs meeting both criteria were calculated and reported as ’Properly paired, quality alignment reads’.

To assess the relationship between alignment quality metrics and sequence variation, additional scatterplots were generated for each sample. For every read pair, the total number of deletions, insertions and single nucleotide polymorphisms (SNPs) was computed, along with the cumulative count of all variant types. These variant counts were thereafter plotted against the corresponding Mapped Identity and Matched Identity scores.

### Chromosomal distribution

Using a custom Python script, the number of mapped read pairs per chromosome, unplaced and unlocalized sequences, respectively, in each sample were plotted.

### Chromosomal mappings

To quantify read coverage across the genome, a custom Python script was developed. The CIGAR string for each aligned read was parsed to extract the reported chromosomal start position, thereafter match operations (“M”) were used to increment coverage counts at the corresponding reference genome coordinate, deletion operations (“D”) were interpreted as gaps in the reference to which no coverage count was calculated and insertion operations (“I”), which represent bases present in the read but not in the reference genome, were accounted for by assigning virtual fractional positions immediately following the aligned reference position, e.g., position *n* were followed by the added position *n*.1 to include the insertion. These fractional positions ensured that insertions were recorded and could be distinguished from reference-based alignments.

The calculated counts per position were plotted, with x-axis depicting genomic position and y-axis the number of reads (count) mapped to each position. In the plot for mitochondrial genome, gene start positions were marked by green horizontal lines and gene end positions by red horizontal lines.

To assess the consistency of read mapping coverage between samples, pairwise Kendall rank correlation coefficients were calculated between pairs of samples using the per-base count values.

### Count filtering

Using a custom Python script, mapped genomic regions were filtered based on read count thresholds to exclude low-coverage areas. After chromosomal mapping, a region was retained only if at least one sample exhibited a minimum count at any position within the region. The minimum threshold value was user-defined (e.g., ≥10 or ≥20 reads) and applied uniformly across all samples.

## Notes

### Competing Interest Statement

The authors have declared no competing interest.

## References

1. Akintunde, O., Tucker, T., & Carabetta, V. J. (2025). The Evolution of Next-Generation Sequencing Technologies. Methods Mol Biol, 2866, 3–29.

2. Andrews, S. (2010). FastQC: A Quality Control Tool for High Throughput Sequence Data [Online]. http://www.bioinformatics.babraham.ac.uk/projects/fastqc/

3. Chon, A., & Huang, X. (2021). Sramm: short read alignment mapping metrics. International Journal of Bioinformatics and Biosciences, 11.

4. Cock, P. J., Fields, C. J., Goto, N., Heuer, M. L., & Rice, P. M. (2010). The Sanger FASTQ file format for sequences with quality scores, and the Solexa/Illumina FASTQ variants. Nucleic Acids Res, 38(6), 1767–1771.

5. Ewels, P., Magnusson, M., Lundin, S., & Kaller, M. (2016). MultiQC: summarize analysis results for multiple tools and samples in a single report. Bioinformatics, 32(19), 3047–3048.

6. Li, H., & Durbin, R. (2010). Fast and accurate long-read alignment with Burrows-Wheeler transform. Bioinformatics, 26(5), 589–595.

7. Li, H., Handsaker, B., Wysoker, A., Fennell, T., Ruan, J., Homer, N., Marth, G., Abecasis, G., Durbin, R., & Genome Project Data Processing, S. (2009). The Sequence Alignment/Map format and SAMtools. Bioinformatics, 25(16), 2078–2079.

8. McCombie, W. R., McPherson, J. D., & Mardis, E. R. (2019). Next-Generation Sequencing Technologies. Cold Spring Harb Perspect Med, 9(11).

9. Reuter, J. A., Spacek, D. V., & Snyder, M. P. (2015). High-throughput sequencing technologies. Mol Cell, 58(4), 586–597.

10. Sherman, M. D., & Mills, R. E. (2018). BAMnostic: an OS-agnostic toolkit for genomic sequence analysis. Journal of Open Source Software, 3(28).

11. Usui, N., Ono, Y., Aramaki, R., Berto, S., Konopka, G., Matsuzaki, H., & Shimada, S. (2021). Early Life Stress Alters Gene Expression and Cytoarchitecture in the Prefrontal Cortex Leading to Social Impairment and Increased Anxiety. Front Genet, 12, 754198.

